# Up-regulated transcriptional regulators in mutant RAS gene signatures: a time-resolved multi-omics study in generic epithelial cell models

**DOI:** 10.1101/2024.06.04.597297

**Authors:** Katharina Kasack, Patrick Metzger, Heiner Koch, Bertram Klinger, Anastasia Malek, Oleg Tchernitsa, Alexander Gross, Wasco Wruck, Balazs Györffy, Bernhard Küster, Christine Sers, Melanie Börries, Reinhold Schäfer

**Affiliations:** Laboratory of Molecular Tumor Pathology, Institute of Pathology and Comprehensive Cancer Center, Charité Universitätsmedizin Berlin, D-10117 Berlin, Germany; Institute of Medical Bioinformatics and Systems Medicine, Medical Center-University of Freiburg, Faculty of Medicine, University of Freiburg, D-79110 Freiburg, Germany; Chair for Proteomics and Bioanalytics, Technical University of Munich, Emil-Erlenmeyer-Forum 5, D-85354 Freising, Germany; Dept. of Bioinformatics, Semmelweis University, H-1094, Budapest; Cancer Biomarker Research Group, Institute of Molecular Life Sciences, HUN-REN Research Centre for Natural Sciences, H-1117, Budapest and Dept. of Biophysics, Medical School, University of Pecs, H-7624, Pecs, Hungary; German Cancer Consortium (DKTK), German Cancer Research Center (DKFZ) Heidelberg, partner sites, Munich; German Cancer Consortium (DKTK), German Cancer Research Center (DKFZ) Heidelberg, partner sites, Berlin; German Cancer Consortium (DKTK), German Cancer Research Center (DKFZ) Heidelberg, partner sites, Freiburg

## Abstract

The expression of mutated RAS genes drives extensive transcriptome alterations. Perturbation experiments have shown that the transcriptional responses to downstream effector pathways are partially unique and non-overlapping, suggesting a modular organization of the RAS-driven expression program. However, the relationship between individual deregulated transcription factors and the entire cancer cell-specific genetic program is poorly understood. To identify potential regulators of the RAS/MAPK-dependent fraction of the genetic program, we monitored transcriptome and proteome changes following conditional, time-resolved expression of mutant HRAS^G12V^ in human epithelial cells during neoplastic conversion. High mobility group AT hook2 (HMGA2), an architectural chromatin modulating protein and oncofetal tumour marker, was recovered as the earliest upregulated transcription factor. Knock-down of HMGA2 reverted anchorage-independent growth and epithelial-mesenchymal transition not only in HRAS-transformed cells but also in an independent, KRAS^G12V^-driven rat epithelial model. Moreover, HMGA2 silencing reverted the deregulated expression of 60% of RAS-responsive target genes. These features qualify HMGA2 as a master regulator of mutant RAS-driven expression patterns. The delayed deregulation of FOSL1, ZEB1 and other transcription factors with known oncogenic activity suggests that HMGA2 acts in concert with a network of regulatory factors to trigger full neoplastic conversion. Although transcription factors are considered difficult to drug, the central role of HMGA2 in the transcription factor network as well as its relevance for cancer prognosis has motivated attempts to block its function using small molecular weight compounds. The further development of direct HMGA2 antagonists may prove useful in cancer cells that have developed resistance to signalling chain inhibition.

## INTRODUCTION

Mutant RAS genes mediate many aspects of malignant cellular transformation (Downward, 2003) and are major drivers of poor clinical prognosis and therapy failure in various types of cancer (Aredo et al., 2019; Bardelli and Siena, 2010; Rui et al., 2015; Vendramini et al., 2022). Mutant RAS proteins act as chronically active signal transducers through downstream RAF/MAPK, PI3K/AKT and RAL effector pathways (Goulding et al., 2020; Martin et al., 2011; Prior et al., 2020; Simanshu et al., 2017). As a consequence of the over-stimulation of cytoplasmic signalling, oncogenic RAS proteins and downstream pathways exert decisive effects on the genetic program of the cells (Zuber et al., 2000). The link between RAS/MAPK/ERK signalling and activation of transcriptional regulators is particularly well documented (Hazzalin and Mahadevan, 2002; Treisman, 1996). MAPK signalling is also involved in chromatin modifications such as histone acetylation and phosphorylation (Yang et al., 2013). Specific transcription factors acting downstream of RAS include members of the ETS protein family (ETS-1, ETS-2) (Plotnik et al., 2014), the bZIP transcription factor family (FOS, JUN) (Bortner et al., 1993), the bHLH protein family (e.g. MYC) (Kapeli and Hurlin, 2011), INTS11 (Yue et al., 2017), Y-box binding protein 1 (Jurchott et al., 2010), KRAB zinc finger proteins (Reggiardo et al., 2022) and Kruppel-like factor 4 (KLF4) (Yang et al., 2013). The number of known nuclear phospho-ERK substrates is constantly increasing along with progress in phosphoproteome analysis (Unal et al., 2017). The functions of MYC (Cole, 1986; Ingvarsson, 1990) and AP1 (FOS/JUN) (Deschamps et al., 1985; Jochum et al., 2001) and other proteins (Ghaleb and Yang, 2017; Lupo et al., 2013) suggest that MAPK-responsive transcription factors not only mediate malignancy-inducing signals, but can also regulate normal growth control and differentiation. Therefore, it remains an open question to what extent these factors are necessary and sufficient for the mutant RAS-mediated malignant phenotype.

Current knowledge about the extent and overall complexity of RAS pathway responsive alterations of the genetic program originated from contrasting expression profiles of phenotypically normal precursor and mutant RAS-transformed cells. Numerous investigations were based e.g. on transcriptome (Huang et al., 2003; Moumtzi et al., 2010; Tchernitsa et al., 2004; Zuber et al., 2000), proteome (Li et al., 2016; Rignall et al., 2009), membrane proteome (Martinko et al., 2018), the secretome (Demory Beckler et al., 2013), glycoproteome (Sudhir et al., 2012) and metabolome analysis (Hutton et al., 2016). In addition, RAS pathway-associated gene signatures were described in cancer cell lines and tumour tissue (Bild et al., 2006; Li et al., 2016; Loboda et al., 2010; Rignall et al., 2009; Stephens et al., 2017; Sweet-Cordero et al., 2005). While these studies identified numerous RAS pathway-responsive targets able to contribute to or to execute malignant phenotypes, the contribution of single or multiple regulators to overarching mechanisms that control the RAS-responsive genetic program remained elusive. Perturbation experiments from our group suggested that RAS pathway-responsive transcription factors function in a hierarchical network and exert distinct phenotypes (Stelniec-Klotz et al., 2012). Furthermore, we found that the overall transcriptional response is restricted by downstream pathway activity. Pharmacological or genetic interference experiments in RAS mutant tumour cells revealed a specific modularity of the transcriptional response, as the expression profiles were largely distinct and overlapped only partially following blockade of the MAPK (Jurchott et al., 2010; Tchernitsa et al., 2004), PI3K (Krech et al., 2010) and RAL (Gyorffy et al., 2015) pathways, respectively.

A major drawback of most expression profiles of cell lines and tumours driven by mutated RAS genes is their snapshot nature, which fails to capture the dynamics of transcription factor-generated processes in cells. Consequently it is challenging to dissect overarching regulatory principles and uncover potential master regulators. To overcome the limitations of static analysis, dynamic approaches that simultaneously assess the interplay between expression profile changes and phenotypic alterations offer significant advantages. Conditional expression systems for mutated RAS genes, which permit time-resolved investigations following transfer into suitable recipient cells, are well-suited for this purpose. The selection of recipient cells is crucial for classifying phenotypic changes, because mutated RAS induces premature senescence in non-immortalized cells with intact tumour suppressor function (Lin et al., 1998; Serrano et al., 1997; Tchernitsa et al., 1999). To recognize neoplastic transformation as the ultimate read-out for mutant RAS expression, we used telomerised, RB- and TP53-deficient human TtH cells, in which conditional RAS activation is driven by HRAS^G12V^ fused to the hormone binding domain of oestrogen receptor (Lim and Counter, 2005). We established a catalogue of RAS-responsive mRNAs, miRNAs, proteins and phosphoproteins. To identify potential regulators of the RAS pathway-mediated cellular expression program, we focused on up-regulated transcription factors. The high mobility group AT hook2 protein (HMGA2) stood out as a very early up- regulated factor. To recognize generic functions and to increase the robustness of our findings, we also screened transcriptome profiles of a mutant KRAS-driven rodent model of ovarian epithelial transformation. Subsequent perturbation of its expression allowed us to identify the architectural transcription factor HMGA2 as a master regulator of oncogene-induced neoplastic transformation.

## RESULTS

### An integrated view of conditional mutant HRAS expression on the genetic program of human epithelial cells

Expression of the oncogene in TtH:RAS:ER cells (Lim and Counter, 2005) was controlled by adding 4-hydroxy-tamoxifen (4-OHT) to the culture medium. We maintained non-induced and induced cells in logarithmic growth phase to exclude cell cycle alterations and effects of cell density following oncogene induction. Prior to HRAS^G12V^ induction, the cells showed the typical epithelial anchorage-dependent phenotype. Expression of the transgene was detectable as early as 4 h, increased between 24 and 72 h, but never exceeded the expression level of the normal allele, unlike other models with forced oncogene overexpression. Phosphorylation of ERK1/2 increased moderately over time. Epithelial-mesenchymal transition (EMT) was visible after 48 h and the cells acquired anchorage independence in semi-solid agar culture (**Fig. 1A-D**). We subjected RNAs and proteins prepared from induced and non-induced TtH:RAS:ER cells isolated at the four time points to RNA-sequencing and mass spectrometry analysis. The overall number of alterations is summarized in **Fig. 1E**. To visualize the time-dependent dynamic responses to HRAS^G12V^ activation, we established self-organizing maps (SOMs), which cluster jointly deregulated gene products as “meta-genes” (**Fig. 2A, Suppl. Table 1A-M**). Major changes of mRNAs and proteins occurred between 4 and 48 h, whereas most dynamic alterations of the phosphoproteome were evident between 4 and 24 h post mutant HRAS induction. The entire catalogue of time-resolved alterations at each molecular level displaying log2-fold changes is shown in **Suppl. Table 2**. To identify associations of mutant HRAS-responsive alterations with biological processes and intracellular networks, we interrogated the ConsensusPathDB interaction database (Herwig et al., 2016) (**Fig. 2B, Suppl. Table 3**). Significant changes of transcriptome and proteome observed as early as 24 hours after mutant HRAS induction involved signalling processes related to IL-10, NFκB, MAPK, WNT, TGF-ß and VEGFR as well as networks of transcriptional regulators such as AP-1, ATF-2 and C-MYB. Some of them showed further upregulation or stabilization 72 hours post oncogene induction, suggesting a role in executing the transformed phenotype. Alterations related to the apoptosis program, cell adhesion processes, signalling processes involving PI3K, ALK1, MTOR and RHO as well to networks including the transcription factors FOXA1, E2F, MYC were exclusively visible at the phosphoprotein level. Underrepresented processes at the proteome and phosphoproteome level involved collagen formation, syndecan signal transduction, microtubule regulation, the TP53 tumour suppressor network, ubiquitin-mediated proteolysis, as well as the protein kinases AKT, Aurora A, and PLK1.

**Fig. 1.**
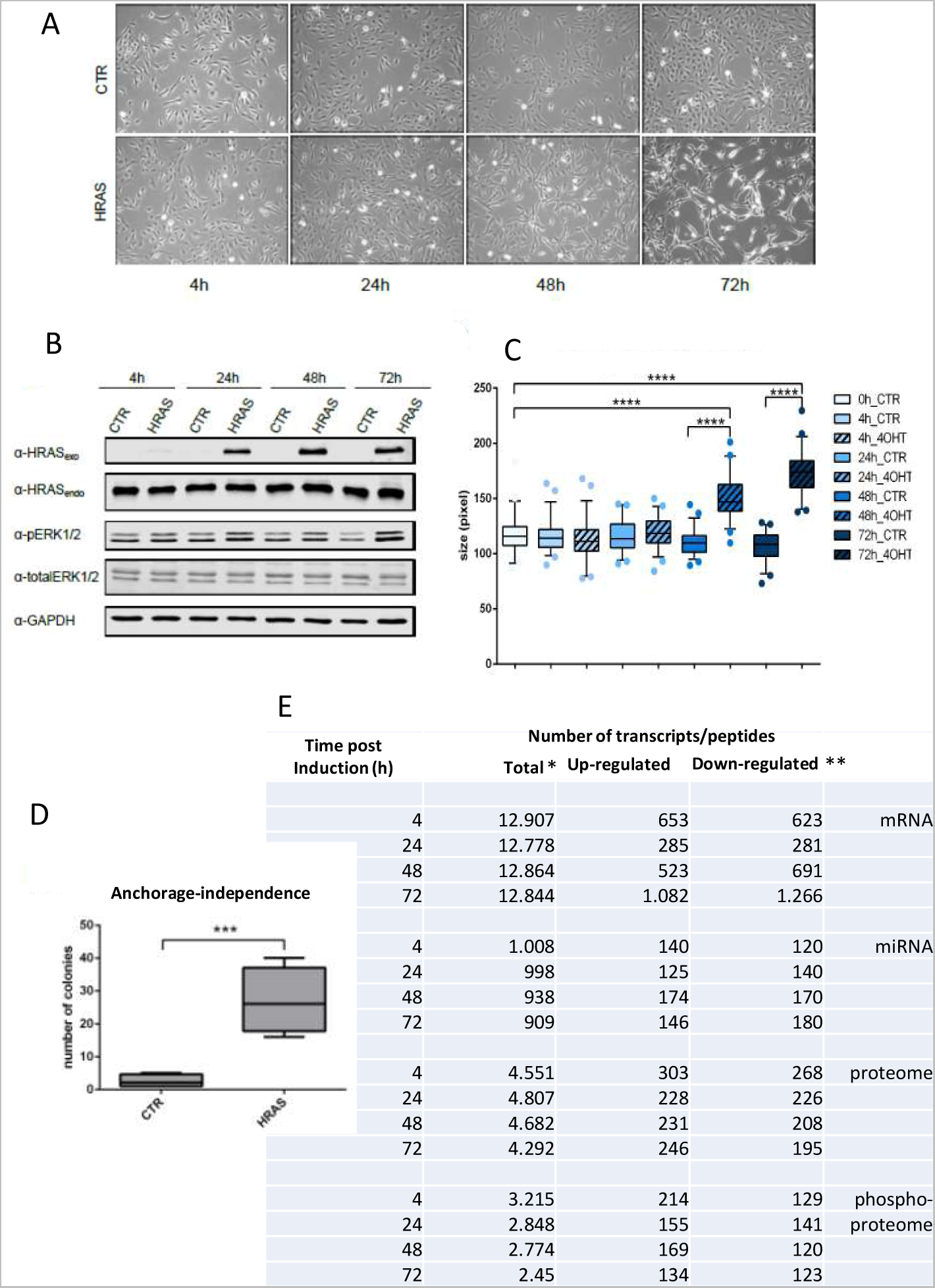
Time-resolved expression of mutant HRAS^G12V^ and phenotypic effects. **A**, Phase contrast images of non-induced, solvent treated (CTR) and 4-OHT-induced TtH ER RAS cells at the indicated time points. Magnification 200-fold. **B**, Western blot analysis of total cell lysates showing protein levels of endogenous HRAS, transgenic HRAS:estrogen receptor hormone binding domain fusion protein, P-ERK1/2, total ERK and GAPDH (loading control). **C**, Quantification of morphological transformation/epithelial-mesenchymal-like transition of 4-OHT-treated cells. The length of cellular protrusions of 50 cells was measured in triplicate per indicated time point, p-values <0.0001. **D**, Colony formation in semi-solid agar medium of cells induced for 48 h. Assays were done in triplicate, p-value 0.001. **E**, Summary of multi-omics analysis. Number of resolved mRNA transcripts, miRNAs and translation products, *Counts >1 based on expression matrix for RNA-seq; **Filtered for de-regulated mutant HRAS-responsive targets, mRNA adj. p-value <0.001, log FC >1, miRNA adj. p-value <0.1; peptides adj. p-value <0.05; phosphopeptides adj. p-value < 0.05.

**Fig. 2.**
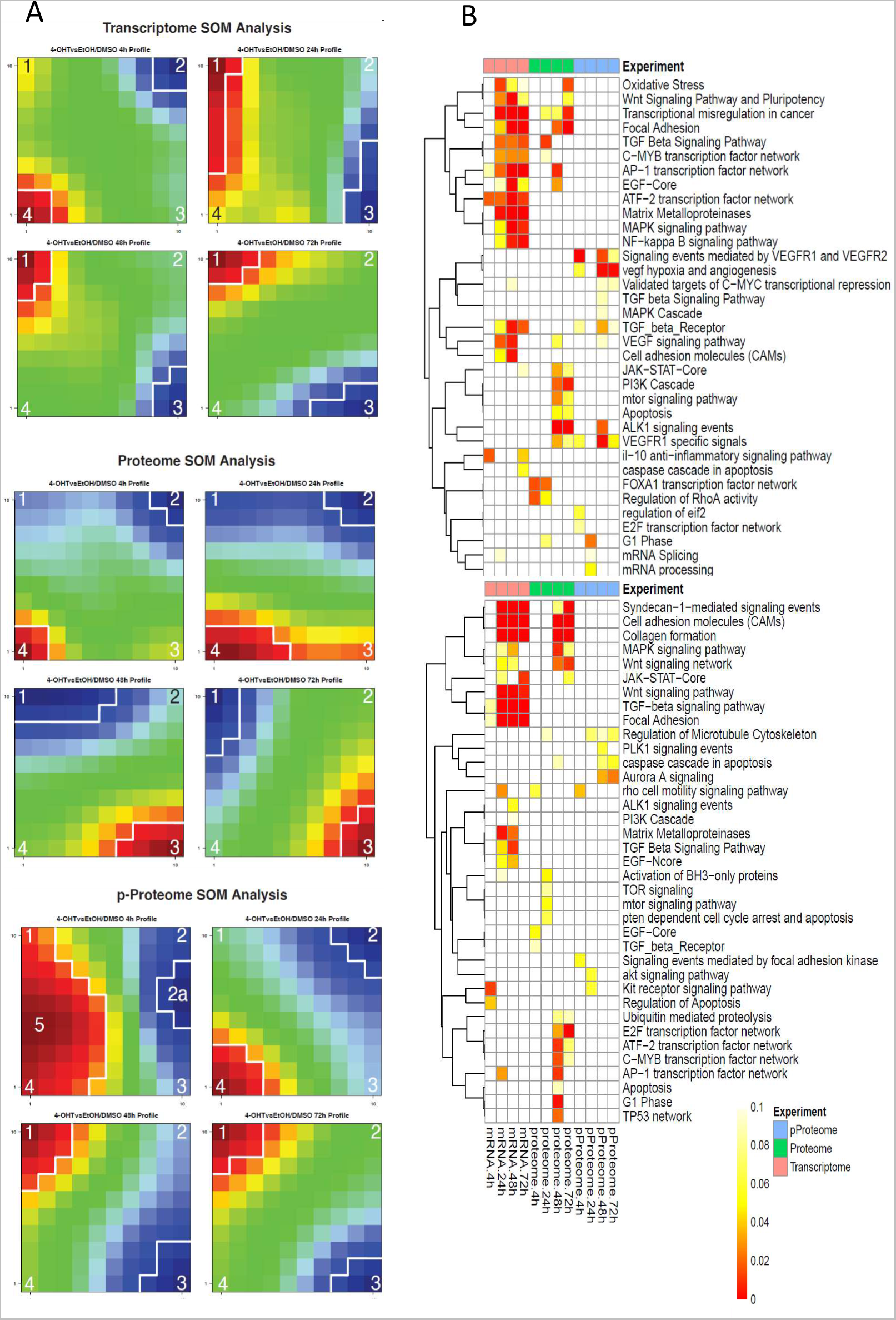
Dynamics of time-resolved alterations of mRNA, protein expression and phosphoprotein levels in TtH:RAS:ER cells and affected biological processes. **A**, Self-organizing maps (SOM) were generated independently for mRNAs, proteins and phosphoproteins using a machine-learning algorithm (Wirth et al., 2011; Wirth et al., 2012). Taking into account their log2-fold expression values, co-regulated transcripts and proteins were clustered into groups designated “meta-genes” or “spots”. Since the number of mutant HRAS-responsive factors varied between approximately 13,000 mRNAs and 500 phosphoproteins, depending on the resolution of the analysis, all responsive targets were fitted into a grid of 10x10 meta-genes. Red spots correspond to groups of up-regulated genes or proteins, taking into account the range from maximally measured expression to 65% of the minimally measured expression, according to previous publications (Wirth et al., 2011; Wirth et al., 2012). Blue spots depict down-regulated genes or proteins considering the range from minimally measured expression to 65% of minimal expression. Within the same class of molecules, the grouped factors are always located at the same (X/Y) position of the grid to enable a comparison between the measured time intervals of 4h, 24h, 48h and 72h. However, the positions are unequal for mRNA, protein and phosphoprotein profiles, because clustering was performed independently. **B**, Biological processes and networks affected by mutant HRAS. Consensus path DB analysis: top, over-represented (enriched); bottom, under-represented pathways and processes.

### Time-resolved upregulation of transcription factors correlates with the process of neoplastic conversion triggered by mutant RAS

Given the large number of RAS-Pathway-dependent changes in the genetic program, the question of overarching regulatory principles of gene regulation in RAS-transformed cells arose. However, it seemed difficult to determine possible general levels of regulation based on single-target gene analyses and, for example, considering the rates of RNA synthesis and degradation, promoter and enhancer structures, chromatin status, or posttranscriptional modifications. Therefore, we used a reductionist approach to search for RAS-responsive transcription factors, annotated in the AnimalDB database (Zhang et al., 2015), which could be considered as master regulators at least for parts of the program, due to their early expression after induction of the oncogene. To increase the robustness of our findings, we focused on transcription factors, whose expression was altered in our screening at the RNA and protein level in a time-resolved manner. We then combined the expression values measured between 4 and 72 hours post HRAS induction into a common data set of 27 deregulated transcription factors (**Fig. 3A**). Next we constructed heat maps representing the RAS pathway responsive, over-expressed and suppressed transcription factors by hierarchical clustering based on Euclidean distance and the complete linkage method (Murtagh, 1985). To verify the two groups, we performed k-means clustering (Hartigan, 1979) (**Fig. 3B**, **Suppl. Table 4**). High mobility group AT hook2 (HMGA2) featured prominently as the earliest up-regulated transcription factor. Moreover, high mobility group AT-hook 1 (HMGA1), v-Maf musculoaponeurotic fibrosarcoma oncogene family protein F (MAFF), FOS-like antigen 1 (FOSL1/FRA-1) and ETS-domain protein 3 (ELK3) formed a second wave of transcription factor upregulation 48 h post RAS induction. Since a clear morphological transformation of the cells was already visible at the end of this time interval (**Fig. 1A**), it is obvious that the upregulation of the transcription factors contributes decisively to the activation of genes necessary for the execution of the neoplastic phenotype. The upregulation of the transcription factor Zinc finger E box-binding homeobox 1 (ZEB1) occurred comparatively late and was correlated with the development of the EMT after 72 hours (**Fig. 1C**).

**Fig. 3.**
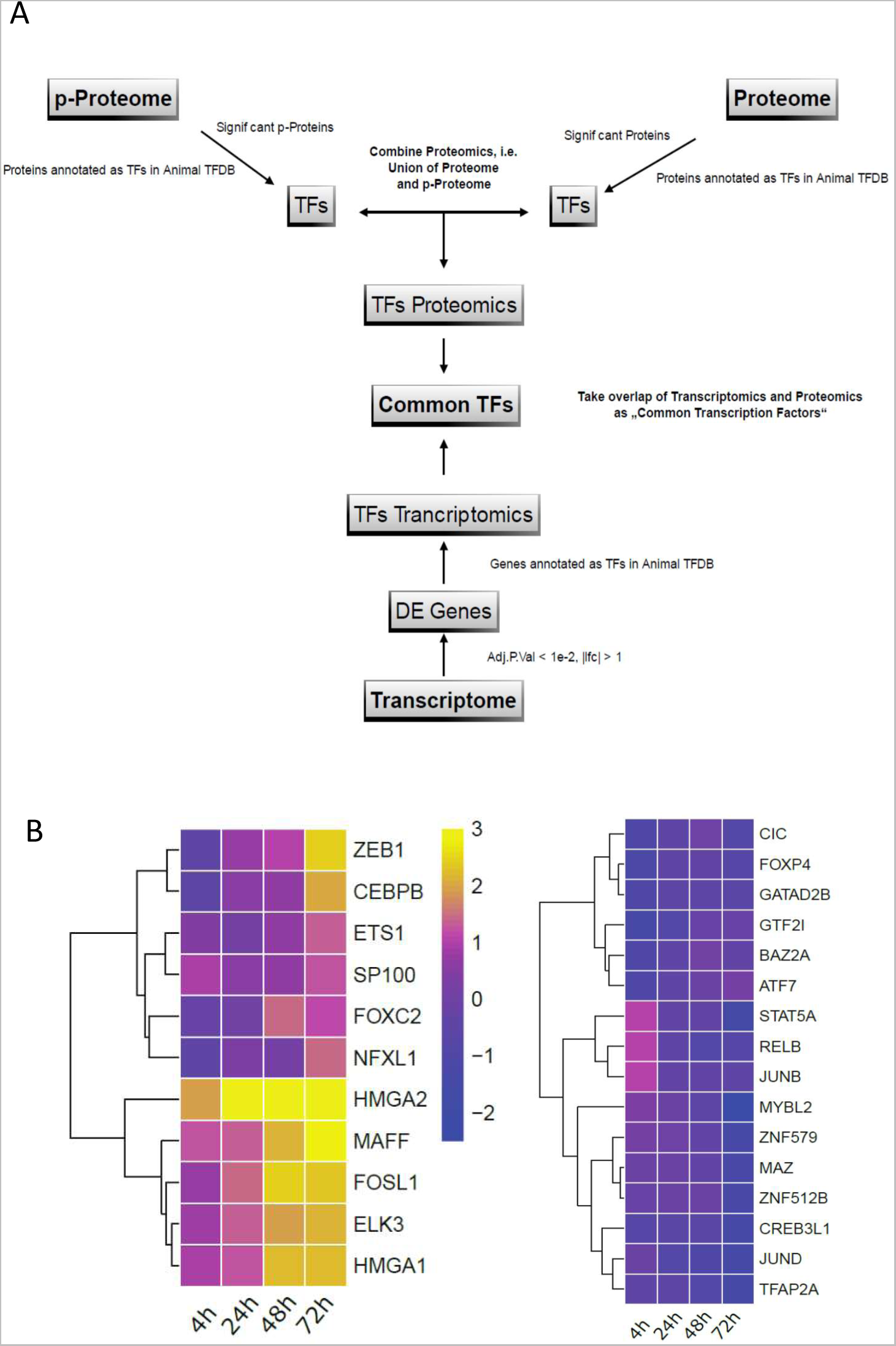
Identification of potential master regulators of HRAS^G12V^-responsive transcriptome. **A,** Strategy to identify robustly deregulated transcription factors in HRAS^G12V^ -responsive transcriptome, proteome and phosphoproteome. Transcription factor annotation based on AnimalTFDB^2.0^. Differentially expressed transcription factor genes and proteins were grouped into a common set. **B**, Heat maps depicting up-regulated (left panel) and down-regulated factors (right panel) following mutant HRAS induction for 4-72h; color code: log2-fold changes. All genes with a FDR (Benjamini, 1995) corrected adjusted p-value smaller than 1e^-2^ and a log2FC > 1 were considered.

### Robust and evolutionary conserved RAS-responsive up-regulation defines architectural factor HMGA2 as a master regulator of the RAS pathway-mediated transcriptome

Following RAS induction gradually causing EMT and anchorage independence, HMGA2 protein expression and phosphorylation on serine 44 increased continuously (**Fig. 4A**). Silencing of HMGA2 expression resulted in reversion of EMT and loss of anchorage independence (**Fig. 4B-D**), indicating that high HMGA2 expression was necessary for the manifestation of the neoplastic phenotype. The RAS/HMGA2 axis was also responsible for neoplastic transformation in the KRAS^G12V^-driven rat surface ovarian epithelial cell model ROSE A2/5 (Stelniec-Klotz et al., 2012; Tchernitsa et al., 2004). In these cells, HMGA2 knockdown resulted in the loss of anchorage independence and in partial reduction of the EMT-like phenotype (**Fig. 4E-G**). The NRAS/HMGA2 axis has also been previously described in rhabdomyosarcoma cells (Li et al., 2013), indicating that the link between RAS signaling and HMGA2 expression is RAS isoform-independent.

**Fig. 4.**
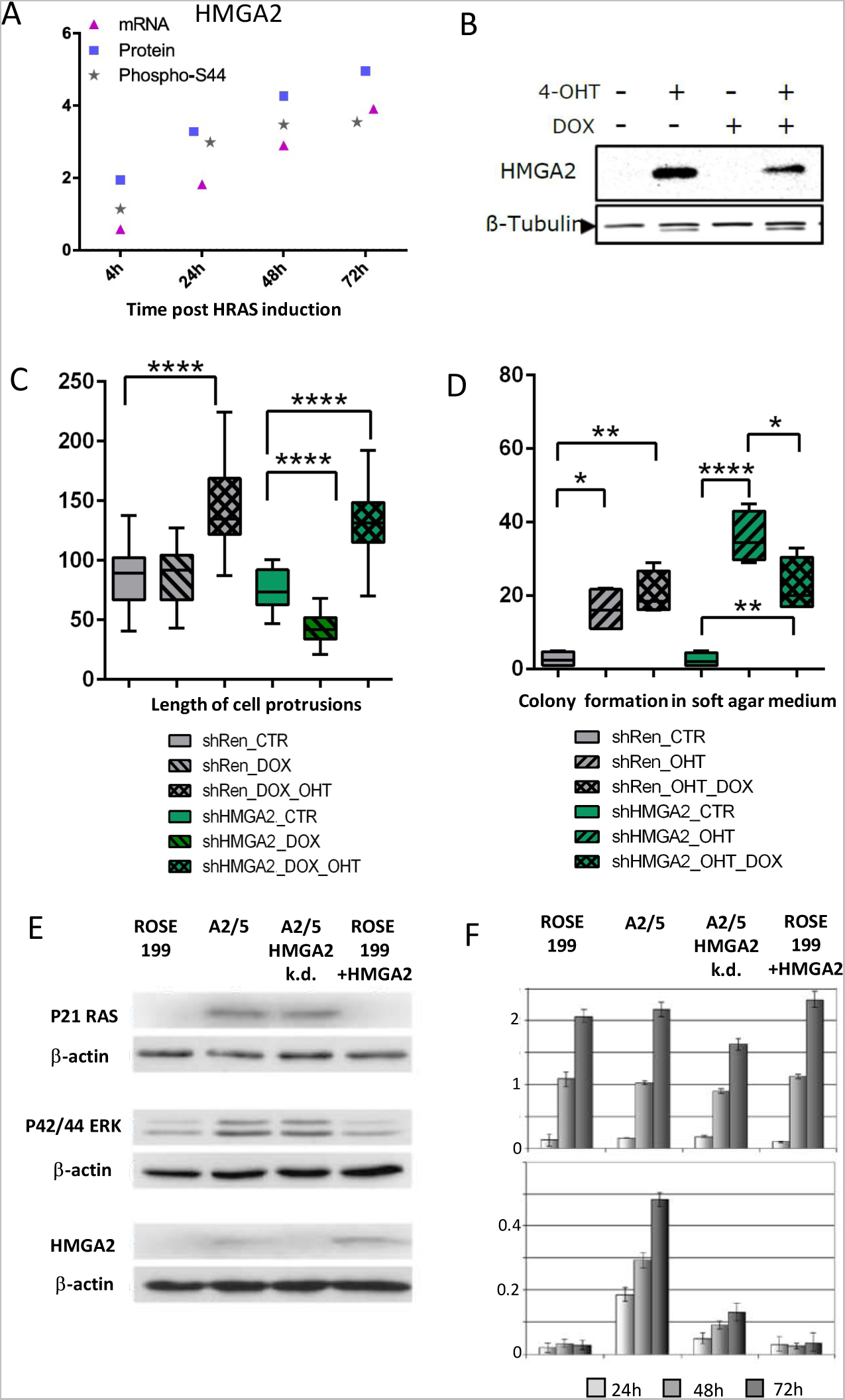
Impact of mutant RAS-responsive HMGA2 expression on cellular phenotypes. **A,** Time course of HMGA2 up-regulation in 4-OHT treated TtH:RAS:ER cells, analyzed at the level of mRNA, protein and phosphorylation at serine 44. **B**, Silencing of HMGA2 expression by introduction of doxycycline-inducible siRNA (shHMGA2) and siRNA targeting Renilla (shRen, control), Western blot analysis, loading control: ß-tubulin. **C**, Effect of HMGA2 silencing on the length of cellular protrusions and **D** on anchorage-independent proliferation. **E**, Western blot analysis of p21RAS, p42/44 Erk, Hmga2 and b-actin expression (loading control) in normal ROSE 199 cells, KRAS^G12V^-transformed ROSE A2/5 cells, A2/5 cells, in which Hmga2 expression was silenced by RNA interference, ROSE 199 cells transfected with an Hmga2 expression vector. **F**, Proliferation in monolayer culture (top panel) and in semi-solid agar medium (bottom panel).

To assess the precise effect of HMGA2 expression on the mutant RAS-responsive transcriptome, we interrogated previously developed customized microarrays for contrasting the expression of 305 validated RAS pathway-dependently regulated transcriptional targets (Tchernitsa et al., 2006; Tchernitsa et al., 2004) in non-transformed progenitor cells ROSE 199 cells (**Fig. 5A**), KRAS-transformed ROSE A2/5 cells (**Fig. 5B**), in ROSE A2/5 cells in which HMGA2 expression was silenced (**Fig. 5C**) and in ROSE 199 cells with forced HMGA2 expression (**Fig. 5D**). The KRAS signature of ROSE A2/5 cells featured 66 up-regulated and 110 down-regulated genes as compared to non-transformed ROSE 199 cells (F**ig. 5E****, Suppl. Table 5**). Silencing of HMGA2 expression neutralized the increased expression of approximately 70% of KRAS^G12V^-responsive, up-regulated genes and reverted the suppression of 60% of down-regulated genes (**Fig. 5F, Suppl. Table 5**). The remaining signature genes only partially reverted to pre-transformation mRNA levels in ROSE 199 cells or their RAS-responsive, deregulated expression was unaffected by HMGA2 knock-down. Remarkably, ectopic HMGA2 expression in ROSE 199 cells neither conferred activation of the MAPK pathway nor induced neoplastic transformation (**Fig. 4F**, **Fig. 5D**). The transcriptional response was distinct form the pattern observed in KRAS-transformed A2/5 cells (**Fig. 5G, Suppl. Table 5**). Altogether, these findings underscored the decisive role of the RAS/MAPK/HMGA2 axis in mediating transformed phenotypes and in shaping a large part of the RAS-responsive transcriptome.

**Fig. 5.**
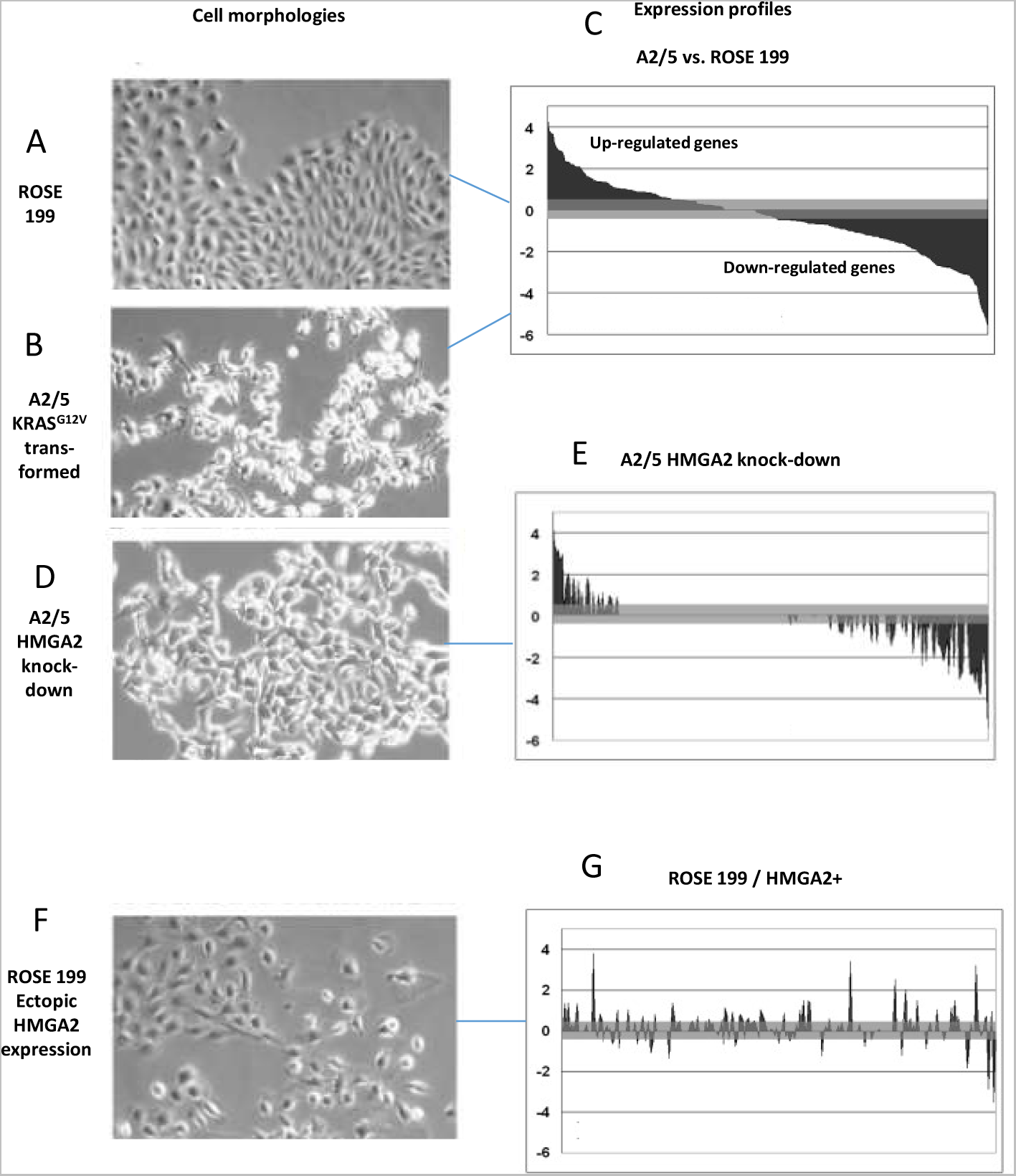
Comparison of phenotypes and transcriptomes of ROSE 199 and KRAS^G12V^-transformed ROSE A2/5 cells. **A**, Morphological characteristics of rat ovarian surface epithelial cells ROSE 199 and **B**, A2/5, KRAS^G12V^- transformed derivatives; **C**, Expression profile showing mutant KRAS mediated upregulation and down-regulation of target genes arranged in descending order according to the log2-fold changes. **D**, A2/5 cells with HMGA2 knock-down and **E**, expression profile of cells arranged as in C; gaps represent genes whose expression reverted to the pattern in ROSE 199. **F**, ROSE 199 cells with forced expression of HMGA2 and **G**, expression profile arranged as above. Expression profiles were generated by interrogation of customized microarrays using dye inversion. Grey areas: non-significant transcriptional alterations. A biological and technical replicate experiment showed 89% reproducibility. Cell images using phase contrast microscopy, magnification 100-fold.

### RAS-responsive alterations of the microRNAome and impact on transcription factor upregulation

In addition to the distinct effects on mRNA expression mutant RAS signaling affects the microRNAome (Anelli et al., 2018; Frezzetti et al., 2011; Kent et al., 2010). The TtH:ER:RAS model also allowed to capture the temporal dynamics of miRNA changes. RAS-responsive miRNAs were transiently up-regulated early and down-regulated later after RAS induction In TtH:RAS:ER cells (e.g. MiR-1-3p), others were lately (MiR-125b-5), gradually downregulated during the observation period between 4 and 72 h (MiR-23-3p) or lately down-regulated (miR-23b-3p, miR-145-5p) (**Suppl. Table 6**). To identify patterns of co-expression or anti-correlated expression between RAS-responsive miRNAs and mRNAs encoding the robustly up-regulated transcription factors in TtH:RAS:ER cells (**Fig. 3B**), we searched the miRTarBase listing confirmed regulatory miRNA:mRNA relationships (Huang et al., 2020). We found as many as 21 miRNAs targeting HMGA2, 19 targeting ZEB1, 18 targeting ETS1, 11 targeting HMGA1, 7 targeting CEBPB, 4 targeting FOSL1 and 2 targeting ELK3 (**Suppl. Table 7**). We used a modified version of the miRlastic R package (Sass et al., 2015) to identify anti-correlated expression of transcription factor mRNAs and their presumptive regulatory miRNAs in the TtH:RAS:ER transcriptome. Elevated expression of HMGA2 was inversely correlated with reduced expression of hsa-miR-196-5p at all 4 time points. Moreover, the HMGA2-targeting hsa-miR-23b-5p, hsa-let-7b(e)-5p and hsa-miR-145-5p were down-regulated between 24 and 72h (**Fig. 6A**, **Suppl. Table 8**). Increased HMGA1 expression correlated with downregulation of hsa-MiR-196b-5p during 4-72h, of hsa-MiR-3916 between 48 and 72h and of hsa-let-7-5p at 72h. RAS-responsive downregulation of hsa-MiR-23b-3p at 72h, hsa-MiR-200b-3p, hsa-MiR-965p, and hsa-MiR-1236-3p between 48 and 72h post oncogene induction correlated with elevated ZEB1 expression. The TtH:RAS:ER expression profiles did not provide evidence for anti-correlated expression between miRNAs and ETS1, FOSL1, ELK3 and CEPB (**Suppl. Table 8**), suggesting that their elevated expression is not modulated by any suppressed miRNA retrieved in the model. A cross-species comparison of human TtH:ER:RAS and rat ROSE A2/5 cells confirmed the robustness of regulatory relationships between miRNAs and HMGA2. The transcriptome of KRAS^G12V^-transformed ROSE A2/5 cells exhibited enhanced HMGA2 expression factor and decreased expression of miR-145-5p, miR-181-5p, miR-let-7b and miR-196b-5p. In addition, stably mutant HRAS-transformed HA1ER cells exhibited the same miRNA pattern (**Fig. 6B, Suppl. Table 9**). In conclusion, the mutant RAS-mediated downregulation of these regulatory miRNAs impresses as an evolutionary conserved mechanism enabling HMGA2 upregulation.

**Fig. 6.**
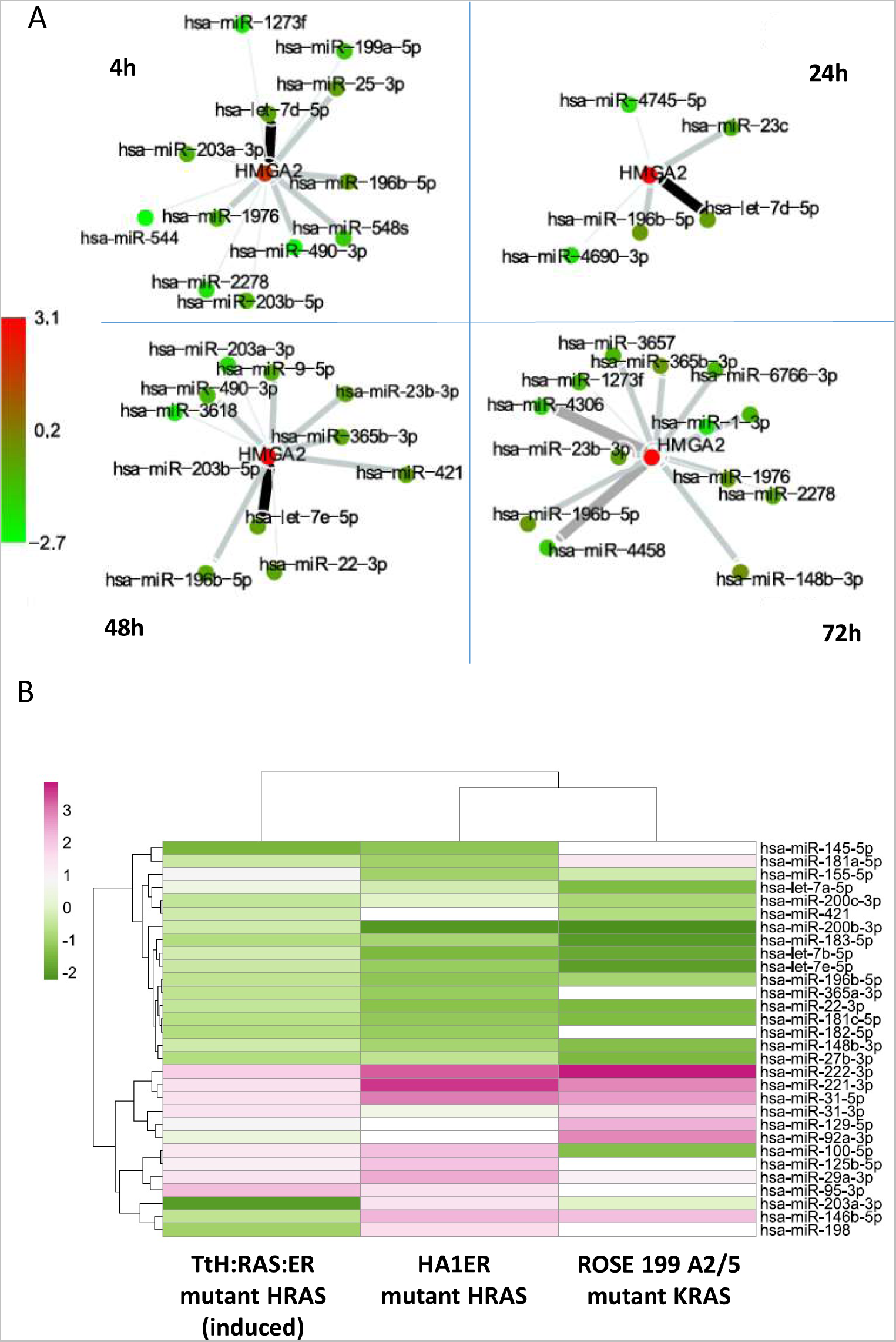
De-regulation of miRNA expression upon conditional or stable mutant RAS expression. **A**, Section of a larger miRNA:mRNA network based on the predicted miRNA targets depicting anti-correlated expression of HMGA2 (up-regulated, red) and putative regulatory miRNAs (down-regulated, green), measured between 4 and 72 h post RAS induction. RNA sequencing data were processed with the miRlastic approach using a linear regression model with elastic net regularization to predict miRNA targets. The elastic net regression was penalized to satisfy the known anti-correlations between miRNAs and mRNAs (Guo et al., 2010). For target prediction, log2fold values were used after contrasting the induced and non-induced TtH:ER:RAS cells at each of the measured time intervals. All miRNAs and mRNAs with an adjusted p-value <0.1 were included. The entire network is not shown. **B**, Heatmap contrasting miRNA expression changes of HRAS^G12V^-induced TtH:RAS:ER cells, stably HRAS^G12V^-transformed HEK cells (HA1ER) and KRAS^G12V^-transformed A2/5 cells. Color code: green – down-regulated, red-up-regulated.

## DISCUSSION

Using an integrated transcriptome and proteome analysis, we investigated transcription factor deregulation in human telomerised, RB and TP53 deficient kidney epithelial cells following conditional HRAS^G12V^ expression over a period of 4-72 hours. Integration of RNA and protein data revealed robust upregulation of high mobility group AT-hook proteins HMGA1 and HMGA2, leucine zipper protein FOSL1 (FRA-1), B-zip transcription factor MAFF, ETS Family member ELK3 (NET) and the EMT transcription factor ZEB-1 during the acquisition of full neoplastic transformation. The most conspicuous upregulation concerned HMGA2, occurring very early compared to the other transcription factors and most pronounced in the period between 24 and 48 hours.

The HMGA2 protein is a small non-histone architectural chromatin binding factor harboring three positively charged AT-hooks as DNA binding domains and a negatively charged C-terminal domain. The AT hooks preferentially recognize AT-rich double-stranded DNA in the minor groove and can DNA segments in *cis* and *trans* (Chen et al., 2010; Cui et al., 2005; Pfannkuche et al., 2009; Reeves and Nissen, 1990). The C-terminal domain is engaged in protein-protein interactions (Gaudreau-Lapierre et al., 2023) and coordinates intramolecular interactions with individual AT-hook domains (Sgarra et al., 2009; Sipe et al., 2022). As a consequence of its architectural function, HMGA2 forms specialized nucleoprotein complexes at enhancers, promoters or branched DNA (Pfannkuche et al., 2009) and invokes heterochromatin structures (Divisato et al., 2022). As a consequence the binding affinities of specific transcription factors to their cognate binding sites can be modulated due to changes in DNA conformation (Cleynen and Van de Ven, 2008; Zhou and Chada, 1998). HMGA2 is expressed in early embryonic and fetal development and in human stem cells, but is silenced in most adult somatic cells (Li et al., 2006; Nishino et al., 2008; Pfannkuche et al., 2009). Typical aberrations in HMGA2 function in cancer are due to overexpression, gene fusion, mutations or truncation (Fusco and Fedele, 2007; Hisaoka et al., 2002; Liu et al., 2020; Young and Narita, 2007). Aberrant HMGA2 re-expression is associated with poor tumor prognosis (Gundlach et al., 2021; Li et al., 2020; Strell et al., 2017; Thi-Hai Pham et al., 2020) and therapy resistance (Dangi-Garimella et al., 2011; Dangi-Garimella et al., 2013; Huldani et al., 2022). Functional studies demonstrated the role of HMGA2 in various aspects of neoplastic transformation (Fedele et al., 2006; Vallone et al., 1997), invasion (Thuault et al., 2006; Watanabe et al., 2009) and metastasis (Wang et al., 2011). However, ablation of HMGA2 and HMGA1 did not reduce the metastatic potential (Chiou et al., 2018), suggesting that other factors acting in a possibly robust network of regulators (Stelniec-Klotz et al., 2012) can compensate for the loss of HMGA function.

Based on the functional classification of growth factor-stimulated genes (Lanahan et al., 1992; Tullai et al., 2007), HMGA2 acts as a typical delayed early response factor. The mode of upregulation and biochemical properties of HMGA2 suggest an important regulatory function in shaping the global transcriptional response to mutational RAS activation. To determine the extent to which HMGA2 upregulation affects the RAS-mediated genetic program, we employed an independent model exhibiting induction and maintenance of neoplastic transformation by stable KRAS^G12V^ expression (Stelniec-Klotz et al., 2012; Tchernitsa et al., 2004). While this model does not capture the early effects of RAS activation, it offers the advantage of easily detecting the effects of HMGA2 ablation on mutant RAS-driven phenotypes and the transcriptome. We interrogated tailored microarrays representing previously characterized and validated RAS pathway transcriptional targets using RNAs from KRAS-transformed A2/5 cells and derivatives, in which HMGA2 expression was knocked-down. Notably, under conditions of HMGA2 ablation 60% of RAS-mediated transcriptional changes reverted almost completely to the pre-transformation level concomitant with the loss of anchorage independence and partial reversion of EMT. The magnitude of transcriptional alterations affecting the RAS gene signature qualifies HMGA2 as a central hub for controlling gene expression associated with transformed phenotypes.

The chromatin remodeling function together with the very early upregulation indicates that HMGA2 plays a superior role in the deregulation of the genetic program in mutant RAS expressing cells. However, to ultimately determine transcriptional patterns and phenotypes HMGA2 is likely to require the concerted action of other transcription factors such as MYC (Yang et al., 2020). This also applies to the co-regulated transcription factors, whose upregulation was delayed after mutant RAS induction. The role of FOSL1 in tumorigenesis is well documented, as is its function as a regulator of transformation-associated genes (Jiang et al., 2020; Milde-Langosch, 2005; Vallejo et al., 2017). FOSL1/FRA1-mediated transformation in a model of thyroid cancer required HMGA2 (Vallone et al., 1997). Silencing of HMGA2 expression reduced FOSL1 expression in ROSE 199 A2/5 cells (this paper). HMGA2 regulates a network of EMT transcription factors including ZEB1 (Tan et al., 2015). Moreover, oncogenic functions were reported for ZEB1 (Krebs et al., 2017), MAFF (Annunziata et al., 2011; Kawai et al., 1992), HMGA1 (Cleynen et al., 2007) and ELK3 (NET) (Semenchenko et al., 2016). The less strongly upregulated transcription factors included CEBPB, known to be required for survival of skin cancer cells under conditions of DNA damage-induced cellular stress (Messenger et al., 2018). The forkhead protein family member FOXC2 triggered EMT and metastasis by activating MAPK and AKT signalling (Cui et al., 2014; Cui et al., 2015; Mani et al., 2007). The involvement of the transcriptional repressor X-box-binding-like-1 (NFXL1) (Song et al., 1994) in RAS-mediated transformation is still unknown, but this factor may be involved in RAS-responsive gene target gene suppression. Interestingly, nuclear antigen SP100 serves as a negative regulator of ETS1 and may limit ETS-mediated transcription (Yordy et al., 2005). This implies that SP100 has the capacity to throttle the activity of effectors and downstream transcriptional targets.

HMGA2 silencing in normal somatic cells is controlled by let-7 (Lee and Dutta, 2007) and other miRNAs (Hashemi et al., 2023). Hence, the down-regulation of let-7 family members by conditional HRAS^G12V^ and stable KRAS^G12V^ expression, respectively, may represent a major, though not exclusive, regulatory mechanism. In TtH:RAS:ER cells, we observed RAS-responsive down-regulation of four microRNAs known to target HMGA2 expression. The pattern of downregulation included early suppression (4h) of miR-196-5p and late suppression of miR-23b-5p, let-7b(e)-5p, and miR-145-5p (24-72 h), suggesting redundant control of HMGA2 via miRNAs. Thus, activation of the RAS/MAPK pathway may simultaneously release these blocks to HMGA2 expression. The functional relationship between miRNAs and HMGA2 expression could be exploited therapeutically (Hashemi et al., 2023)

The increased expression of HMGA2 appears to define a vulnerability of mutant RAS-driven tumor cells. This has motivated multiple approaches to limit HMGA expression in tumors (De Martino et al., 2019). A number of small molecules such as ciclopirox (Huang et al., 2019), niclosamide (Leung et al., 2019) and trabectedin (D’Angelo et al., 2013) may serve as HMGA antagonists. Moreover, other reports have suggested that JQ1 (Kumar et al., 2015), panobinostat, selumetinb (Wachter et al., 2018) and phenformin (Jiang et al., 2016) can suppress HMGA2 expression in tumour cell lines. The described inhibitory effects are likely to be due to indirect inhibition of HMGA2 function. The histone deacetylase inhibitor panobinostat and the bromodomain protein inhibitor JQ1 are known to provoke multiple inhibitory effects. For example JQ1 treatment simultaneously suppressed FOSL1, c-MYC and HMGA2 expression (Kumar et al., 2015). In a recent attempt to identify direct HMGA2 antagonists, we have shown that the FGFR inhibitor PD173094 directly binds to the C-terminus of the protein and modulates its DNA binding and transcriptional activation functions (Ahmed et al., 2023).

In conclusion, the prognostic relevance in cancer and phenotypic functions of HMGA2 warrant further testing of candidate antagonists in mutant RAS-driven tumor cells. Suitable compounds that phenocopy the effects of genetic HMGA2 ablation through inhibiting anchorage independence and EMT rather than normal proliferation might be of particular therapeutic significance. HMGA2 antagonists might enhance inhibitory effects due to signaling pathway blockade by kinase inhibitors (Papke and Der, 2017) or novel RAS inhibitors, especially under conditions of rapidly evolving therapy resistance (Adachi et al., 2020; Awad et al., 2021; Tanaka et al., 2021). We identified HMGA2 as a pivotal early regulator of the mutant RAS-driven genetic program in that its ablation reverts deregulated expression of the majority of RAS-responsive transcriptional targets. This positions HMGA2 as a master regulator of mutant RAS-driven expression patterns. Perturbation experiments targeting the co-regulated transcription factors will further elucidate the entire network of regulatory factors that ultimately orchestrate full neoplastic transformation and even therapy resistance.

## Acknowledgements

Our work was supported by Deutsche Krebshilfe (grant 10-2187-Schä 2 to R.S., grant 70114307 to B.K.), Deutsche Forschungsgemeinschaft (SFB 618, grants to R.S., C.S.), Berliner Krebsgesellschaft (Ernst-von-Leyden fellowship to A.M.) and the German Cancer Consortium (DKTK). We thank A. Geflitter for excellent technical assistance.

## Supplementary information

https://zenodo.org/doi/10.5281/zenodo.11346461

**Suppl. Table 1A-M**

Original data showing the mRNAs, proteins and phosphoproteins clustered in metagenes for 4 time points post RAS induction. A-D mRNAs at 4, 24, 48, 92 h; E-H proteins at 4, 24, 48, 92 h; I-M Phospho-proteins at 4, 24, 48, 92 h

**Suppl. Table 2**

Master Table: Time resolved alterations mRNAs, miRNAs, proteins and phosphoproteins; log2fold changes 4h, 24h, 48h and 72h post induction of TtH ER-Ras cells. Alterations of phosphoproteins and miRNAs are shown in separate spreadsheets named “Phosphoproteins” and “microRNAs”; NA = not detected; Y = values exist in separate table sheets

**Suppl. Table 3**

Overview: Impact of RAS induction on biological processed and pathways (consensus PathDB analysis)

**Suppl. Table 4**

Differentially expressed RAS-responsive transcription factor mRNAs at 4 time points and solvent-treated controls

**Suppl. Table 5**

Effects of HMGA2 silencing on KRAS signature in ROSE199 FINAL

**Suppl. Table 6**

Differentially expressed RAS-responsive miRNAs in TtH:ER:RAS cells

**Suppl. Table 7**

Summary of miRTarBase information listing confirmed regulatory miRNA:mRNA relationships. Excel spreadsheets designated “all” shows all targeting miRNAs, regardless whether regulatory relationships were detected by reported assay, western blot, pPCR, microarray analysis, pSilac or CLIP-Seq. Filtered spreadsheets designated “sorted” show targeting miRNAs, where the impact on mRNA expression was validated throughout by reporter assays, western blots and qPCR.

**Suppl. Table 8**

Summary of down-regulated miRNAs in TtH:RAS:ER cells validated by RNA-sequencing and strength of the miRTarBase prediction for targeting HMGA2, HMGA1, ZEB1, ELK3, MAFF and ETS2. The beta (b) value indicates the strength of the prediction, the lower the value the better the prediction. We used a cut-off of b =0.3.

**Suppl. Table 9**

RAS-responsive miRNA deregulation in TtH:RAS:ER cells, stable mutant HRAS transformants HA1ER and mutant KRAS-transformed ROSE 199 vells A2/5. Comparison of miRNA expression **in** stable HRAS^G12V^-transformed HEK cells, KRAS^G12V^-transformed ROSE A2/5 and induced TtH:RAS:ER cells. Microarray analysis, heatmap, green: downregulated; Red: up-regulated.

Accession numbers of microarray data

GSE 14658. https://www.ncbi.nlm.nih.gov/geo/query/acc.cgi?acc=GSE14658.

GSE46794 (https://www.ncbi.nlm.nih.gov/geo/query/acc.cgi?acc=GSE46794)

GSE46301 (https://www.ncbi.nlm.nih.gov/geo/query/acc.cgi?acc=GSE46301)

## MATERIALS AND METHODS

### Cell lines and cell culture

TtH:RAS:ER cells (obtained from Channing J. Der’s lab, UNC at Chapel Hill) were cultured in low glucose, phenol red-free Dulbecco’s Modified Eagle’s Medium (DMEM) supplemented with 10% fetal calf serum (FCS), 2 mM L-glutamine, penicillin (100U/ml) and streptomycin (100 μg/ml) (standard medium) at 37°C and 5% CO_2_ in a humidified incubator. Expression of HRAS^G12V^ was induced by adding 4-Hydroxytamoxifen (4-OHT) (20nM). Preneoplastic ROSE 199 and KRAS^G12V^-transformed ROSE A2/5 cells (Tchernitsa et al., 2004) were cultured in phenol-red containing DMEM supplemented with 10% FCS and the other additives. ROSE 199 cells were transfected with the KRAS^G12V^ gene cloned under the control of the elongation factor promoter into the expression vector pEF-BOS. To generate ROSE 199 cultures ectopically expressing HMGA2, cells were transfected with the full-length HMGA2 gene cloned under the control of the elongation factor promoter into the expression vector pEF6 (Invitrogen). Stable transfectants were selected in medium supplemented with 10 μg/ml blasticidin. HRAS^G12V^-transformed human embryonic kidney cells (HA1ER) and their preneoplastic precursor line HA1EB (obtained from Robert A Weinberg’s lab, Whitehead Institute) (Hahn et al., 1999) were cultured in Minimum Essential Eagle’s medium, alpha modification (Sigma), supplemented with 10% FCS, 2 mM ultraglutamine, penicillin/streptomycin, 0.1 mg/ml hygromycin B and 0.5 μg/ml puromycin. All cell lines were authenticated by short tandem repeat (STR) analysis (CLC Cell Line Service GmbH) and routinely checked free of mycoplasma contamination.

### Proliferation assays, monitoring of anchorage independence and morphological transformation

To determine cellular proliferation in 96-well cultures we used the AlamarBlue reagent (TREK Diagnostic Systems) or the sodium 3’ [1-(phenyl-amino-carbonyl)-3, 4 tetrazolium]-bis(4-methoxy-6-nitro)-benzene sulfonic acid hydrate (XTT)-based colorimetric assay (Roche) according to manufacturer-recommended protocols. Anchorage-independent growth was assessed in 96 well plates coated with poly-2-hydroxyethyl methacrylate (poly-HEMA) (Sigma). To coat the wells prior to seeding of cells, we added a 5 μg/ml poly-HEMA stock solution (75 µl) dissolved in 95% ethanol to each individual well and dried the plates for 72 h at 37 °C. All assays were performed in triplicate and independently repeated at least once. Colony assays were done in T25 cell culture flasks in standard medium supplemented with semi-solid 0.15% Difco Noble agar. The agar (0.3%) was autoclaved and after cooling-down maintained at 45°C. Cell suspensions (1 ml containing the desired cell number) were pipetted into double concentrated standard medium (32 ml) and the same volume of agar was quickly added. After careful overhead shaking the flasks were placed in an ice bath for a short time to cool and harden the agar. This was followed by incubation at 37°C. Colonies were counted after 14 days. The length of cellular protrusions following induction of HRAS^G12V^ in TtH:RAS:ER cells was measured by microscopy using a Keyence-all-in-One fluorescence microscope and built-in software.

### RNA-interference experiments

*ROSE A2/5 cells*: Deprotected and desalted RNA oligonucleotides obtained from Dharmacon (Lafayette) were used for preparation of RNA duplexes. For annealing of siRNAs, 20 µM single-stranded 21-23 nt RNAs were incubated in annealing buffer (100 mM potassium acetate, 30 mM HEPES-KOH, pH 7.4, 2 mM magnesium acetate) for 1 min at 90 °C and cooled down to room temperature. Sequence of RNA-duplex targeting rat HMGA2: 5’-AACUCCCGAGCCGUAGCGGdTdT-3’ (sense strand) and 5’-CCGCUACGGCUCGGGAGUUdTdT-3’ (anti-sense strand). Sequence of scrambled RNA-duplex for control experiments: 5’-AACUGCGCAGCCGGAUCGGdTdT-3 (sense), 5’-CCGAUCCGGCUGCGCAGUUdTdT-3’ (anti-sense). At 24 h prior to transfection, 4×10^4^ ROSE-A2/5 cells were seeded into 6-well plates. Transfection of siRNA was carried out using Oligofectamine (Life Technologies) and 0.2 μM siRNA duplex. Cells were transfected with siRNA twice in 24 h-intervals and analysed 48 h after the first treatment. We also treated cells with transfection reagents only to provide an additional control besides transfecting scrambled siRNA duplexes.

*TtH:RAS:ER* cells: For knock-down experiments in TtH:RAS:ER cells we generated transfectants expressing modified derivatives of the doxycycline-inducible 00425_pSIN-TRE3G-turboGFP-miRE-PGK-Neo vector (obtained from Johannes Zuber’s lab, Vienna), harbouring 97-mer guide and flanking oligonucleotides targeting HMGA2 and controls (purchased from Integrated DNA Technologies). Oligonucleotides were designed by a proprietary algorithm (Fellmann et al., 2011). Control cells harboured 00425_pSIN-TRE3G-turboGFP-miRE-scrbl-PGK-Neo, expressing scrambled shRNA or (p425) 00425_pSIN-TRE3G-turboGFP-miRE-Ren.713-PGK-Neo, expressing shRNA targeting Renilla, (p245)-00425_pSIN-TREG3-turboGFP-miRE-HMGA2.988-PGK-Neo or (p245)-00425_pSIN-TREG3-turboGFP-miRE-HMGA2.998-PGK-Neo, targeting HMGA2. The following oligonucleotide sequences were used for HMGA2 knock-down (guide oligo italicized):

HMGA2.988:

TGCTGTTGACAGTGAGCGATCCCTCTAAAGCAGCTCAAAATAGTGAAGCCACAGATGTA*TTTTGAGCTGC TTTAGAGGGAC*TGCCTACTGCCTCGGA;

HMGA2.998:

TGCTGTTGACAGTGAGCGCGCAGCTCAAAAGAAAGCAGAATAGTGAAGCCACAGATGTAT*TCTGCTTTCT TTTGAGCTGCT*TGCCTACTGCCTCGGA.

Silencing was induced by addition of 0.25 μg/ml doxycycline to the culture medium. Prior to induction of RAS with 4-OHT and shRNA expression with doxycycline, we seeded 1×10^5^ cells in 6-well plates. Cells were inspected under the microscopy and harvested 48 h after shRNA expression.

### Western blot analysis

Cells were washed twice with cold PBS buffer. To analyze RAS protein levels, membrane protein fractions were extracted with hypotonic lysis buffer containing Tris-HCL 10mM, PMSF 1mM, and proteinase inhibitor, pH8 (Roche). Whole cellular extracts for phospho-p44/42 ERK detection were prepared by lysis in a buffer prepared from the following solutions (solution 1 [9 volumes]: Tris-HCl 10mM, NaCl 150 mM, TritonX-100 1%, DOC 1% pH 7,2; solution 2 [2 volumes]: PMSF 1 mM, NaF 50 mM, leupeptin 50 μg/ml, Na-orthovanadate 1 mM, Aprotinin 4 μg/ml). For HMGA2 protein analysis, cells were subjected to two repeated freeze-thaw cycles and afterwards lysed in the following buffer: 10 mM Hepes (pH7.9), 400 mM NaCl, 0.1 mM EGTA, 5% glycerol, 0.5 mM DTT, 0.5 mM PMSF. The protein extracts were separated by SDS-PAGE and transferred to polyvinyl membranes (Amersham Pharmacia). The following antibodies were used: rabbit anti-phospho-p44/42 MAPK (Thr202/Tyr204) (Cell Signaling, Technology), mouse anti-pan Ras (Transduction Laboratories), rabbit anti HRAS (Proteintech), anti-HMGI-C (Eurogentec), rabbit anti-human HMGA2 (Cell Signaling) or rabbit anti human HMGA2 (New England Biolabs),. Blots were developed using appropriate secondary antibodies and the ECL system (Amersham Pharmacia). To confirm equal protein loading, blots were stripped (Western Blot recycling kit, Alpha Diagnostics, San Antonio, TX, USA) and re-probed with an actin antibody (Chemicon), Gapdh antibody (Santa Cruz Biotechnology) or ß-tubulin antibody (Cell Signaling Technology).

### Transcriptome analysis / RNA Sequencing

TtH:RAS:ER cells plated in 10 cm dishes were induced by addition of 4-OHT for 4, 14, 48 and 72h. Total RNA and miRNA were isolated using the TRIZOL™ reagent according to the manufacturer’s instructions (Invitrogen). RNA library preparation and sequencing was performed at the Genomics and Proteomics Core Facility at the DKFZ in Heidelberg. The TruSeq-Kit v2 (Illumina) was used for library preparation of total RNAs and Next small RNA Kit (New England Biolabs) for library preparation of microRNAs according to manufacturerś instructions. Sequencing was performed on an Illumina HiSeq 2000 v3 with 50 bp single-reads. Quality control of raw reads was done using FastQC analysis (Version 0.10.0).

Sequencing data obtained from RNA and microRNA samples were normalized and filtered with limma (Ritchie et al., 2015). Differentially expressed genes (DEG) and differentially expressed microRNAs (DEM) were identified by calculating the differences between the 4-OHT and the EtOH/DMSO treated control group separately for each of the four time points. DEGs and DEMs were sorted according to their FDR adjusted p-value (Benjamini, 1995). DEGs were reported if the adjusted p-value was <0.01 and the log2FC was >1. DEMs were reported if the adjusted p-value was < 0.1. For each comparison, a functional enrichment (hyperG Test) was performed based on Consensus Pathways (Herwig et al., 2016).

### Transcriptome analysis / Customized microarrays

The customized microarray for detection of differential gene expression in mutant RAS transformed rodent cells was designed as described (Tchernitsa et al., 2006). Briefly, we selected 60-mer oligonucleotides for interrogating genes recovered from subtracted cDNA libraries representing sequences differentially expressed in parental and mutant RAS–transformed cells (Tchernitsa et al., 2004; Zuber et al., 2000). Thirteen genes described as housekeeping genes in many different tissues and cell lines were included to permit normalization of hybridization intensities. The specificity of all probes assembled on the microarray was independently confirmed by analysing the same target RNAs by reverse and conventional northern blotting or RT-PCR. The total number of gene probes is 324 on the array. Total RNA from ROSE 199 cells and derivatives was prepared under conditions of sub-confluency with the TRIZOL reagent (Invitrogen) according to the manufacturer’s instructions.

We applied dendrimer technology for cDNA labelling and array hybridization using the 3DNA Expression Array Detection Kit for Microarray (Genisphere). Two samples of cDNA were synthesized from each RNA probe (20 μg) by reverse transcription with oligo-(dT) primers containing capture sequences complimentary to Cy3-zor Cy5-fluorescent dyes. The cDNA samples carrying the appropriate capture sequences were mixed in the following combinations: cDNAs from ROSE 199 cells with KRAS-transformed ROSE 199 A2/5 cells as well as ROSE 199 cells overexpressing HMGA2 with ROSE 199 A2/5 cells following siRNA-mediated HMGA2 knockdown. In control hybridizations, we mixed ROSE 199 cDNA with cDNA from empty vector transfected derivatives and ROSE-199 A2/5 cells transfected with scrambled siRNA. The concentration of cDNA, hybridization and washing were carried out according to the kit manual. The second round of hybridization was performed in a reaction mix containing equal amounts of 3DNA capture Reagent Cy3 and 3DNA Capture Reagent Cy5. The composition of the reaction mix, hybridization and washing conditions were as described in the Genisphere instruction manual.

Microarray images were obtained on an Agilent G2565AA scanner at 10 μm resolution. Imagene software (BioDiscovery) was used for feature extraction. The significance of expression changes between pairs of cells were evaluated by SAM analysis (Tusher et al., 2001). The mean values of spotted replica on the microarrays were processed further. Lowess intensity dependent normalization was used to adjust for differences in labelling intensities of the Cy3 and Cy5 dyes (Berger et al., 2004). Analyses were performed using BRB-Array Tools (Version 3.7.0) (Simon et al., 2007).

#### miRNA microarray design and hybridisation

Total RNA was isolated from cell lines using the mirVanaTM miRNA Isolation Kit from Ambion® according to the manufacturer’s instructions. In order to avoid cross-labelling of premature miRNA we used the gel purification option. Subsequently the quality of the small RNA fraction was controlled on a 15% PAAG gel stained with ethidium bromide. MiRNA microarrays were constructed using the NCODETM multi-species microarray probe set V1 (Invitrogen). Oligos were spotted at a concentration of 30µM on epoxy treated slides (Corning®) using the MicroGrid compact microarrayer (Genomic Solutions) equipped with a MicroSpot 2500 quill pins. 100 ng of small RNA fraction were labelled using the NCODETM miRNA labelling system (Invitrogen) according to the manufacturer’s instructions. Microarray hybridisation and washing was performed according to the same manual on Slide Booster^TM^ hybridisation station (Advalytix). Ambion miRNA microarrays were hybridized according to provider’s specifications. Microarray images were obtained on an Agilent G2565AA scanner at 10 µm resolution. Image analysis was carried out using the ImaGene software (BioDiscovery). For each target miRNA, two independent hybridizations were done by inverting the Alexa3 and Alexa5 fluorochromes in the labelling reactions. Background subtraction and normalisation was done with GeneSite software (BioDiscovery). For statistical analysis BRB-ArrayTools software was used. The repeated measurements of the duplicated spots on the arrays showed a high reproducibility (Pearson correlation coefficient = 0.94). Filtering was carried out to exclude genes having intensity less than (twice of rounded background average). Normalization was done by centring each array using median over entire array, using the median as reference array. Dye swap experiments were used to control normalization. To select miRNAs differentially expressed between paired normal and transformed samples we used Statistical Analysis of the microarrays (SAM). The significance for SAM was set at 10% of FDR with at list 1.5 fold regulation.

### Proteome and Phosphoproteome analysis

#### Cell lysis

Induced and control TtH:RAS:ER cells grown under conditions of sub-confluency at each time point (as for the transcriptome analysis) were washed two times with PBS and lysed by scraping into 500 μl per plate of lysis buffer (40 mM Tris/HCl pH 7.6, 8 M urea, EDTA-free protease inhibitor complete mini [Roche], and phosphatase inhibitor cocktails 1, 2, and 3 [Sigma-Aldrich] at 1× final concentration according to the manufacturer’s instructions) and a 10 min incubation on ice. The lysate was transferred into reaction vessels and homogenized for 1 min using a Bioruptor® (Diagenode) set to medium power for 1 min at 4 °C. The protein concentration was determined with a Bradford assay (Pierce).

#### Protein digestion, peptide purification and dimethyl labelling

A total of 1,800 µg of protein per experimental condition was used for digestion. Disulfide bonds were reduced with DTT at a final concentration of 10 µM in a thermo shaker at 37 °C and 700 rpm for 60 min. Cysteine residues were alkylated using 55mM chloroacetamide for 30 min at room temperature in the dark. The sample was diluted with three volumes of 40 mM Tris/HCl pH 7.6 to decrease the urea concentration to 1.5 M. Trypsin was added to the cell lysate at a protease-to-protein ratio of of 1:50 (w/w), followed by digestion in a thermo shaker at 37 °C and 700 rpm for 4 h. A second aliquot of trypsin was added at the same protease-to-protein ratio and incubation was contunued at 37 °C overnight. The following day, the samples were cooled to room temperature and acidified using 0.5% TFA. Following the precipitation of insoluble debris at 5000 x g, the supernatant was desalted using 50 mg Sep-Pak columns (Waters) and a vacuum manifold. Columns were primed with 2x1 ml of solvent B (0.07% TFA, 50% ACN) and equilibrated with 3x1 ml of solvent A (0.07% TFA in deionized water). The sample was then slowly passed through the column to allow proper peptide binding. Peptides were washed five times with 1 ml of solvent A and labelling was performed five times with 1 mL of the appropriate labelling reagent. Light labelling reagent per sample: 4 ml of 50 mM sodium phosphate buffer, 500 µl of 600 mM NaBH3CN and 500 µl of 4% Formaldehyde; intermediate labelling reagent per sample: 4 ml of 50 mM sodium phosphate buffer, 500 µl of 600 mM NaBH3CN and 500 µl of 4% D-formaldehyde; heavy labelling reagent per sample: 4 ml of 50 mM sodium phosphate buffer, 500 µl of 600 mM NaBD3CN and 500 µl of D-^13^C-formaldehyde. Peptides were eluted into a reaction vessel using 2 times 150 μl of solvent B. Finally, the samples were frozen at −80 °C and dried completely in a speed vac.

#### Phosphopeptide enrichment using Fe-IMAC columns

Enrichment was essentially performed as previously described (Ruprecht et al., 2015). Briefly, a Fe-IMAC column (ProPac IMAC-10 column, 4 x 50 mm, Thermo Scientific) was charged with Fe^3+^ ions using 3 ml FeCl3 (Sigma Aldrich). Next, the column was equilibrated with solvent A (30 % ACN, 0.07 % TFA) and connected to an Aekta FPLC system (GE Healthcare). Subsequently, 3 mg dimethyl-labelled peptides were reconstituted in 500 µl solvent A and loaded onto the column (5 min, 0.2 ml/min). Peptides were eluted with solvent B (0.3 % NH4OH) in a stepwise gradient of 0-16 % (5-6.72 min, 3 ml/min), 16-26.25 % (6.7-11.7 min, 0.55 ml/min), 26.25-50 % (11.7-12.35 min, 3 ml/min), and 12.35-15 min 0 % (3 ml/min). The phosphopeptide-containing fraction (UV) was collected in a reaction vessel (1 ml) and was evaporated to dryness in a speed vac.

#### High pH reversed-phase micro-column fractionation for phosphopeptides

High pH reversed-phase micro-columns were prepared in 200 µl pipette tips. Five discs (Ø 1.5 mm) of C18 material (3 M Empore) per micro-column were squeezed into a 200 µl pipette tip that was fixed in a 1.5 ml reaction vessel. All solvents were passed through the columns by centrifugation. Columns were primed with 40 µl 50 % ACN, 25 mM NH4COOH (pH 10) and equilibrated with two times 40 µl 25 mM NH4COOH (pH 10). The sample was then slowly passed through the column and the flow-through was collected and applied to a low pH micro-column for desalting as described previously (Rappsilber et al., 2007). Peptides were fractionated using solvents with increasing ACN concentrations (5 %, 7.5 %, 10 %, 12.5 %, 15 %, 17.5 % and 50% ACN in 25 mM NH4COOH, pH10 respectively). The desalted flow-through was combined with the 17.5 % fraction and the 50 % fraction was combined with the 5 % fraction, leading to a total of six fractions. The fractionated samples were dried down prior to mass spectrometric analysis.

#### Hydrophilic strong anion exchange chromatography (hSAX) for peptide fractionation

For full proteome analysis, peptides were fractionated using hydroxide-selective anion-exchange chromatography as described previously (Ritorto et al., 2013). Briefly, an IonPac AS24 analytical column and IonPac AG24 guard column (2 x 250 mm and 2 x 50 mm, Thermo Scientific) were connected to a Dionex Ultimate 3000 HPLC system (Thermo Scientific). An amount of 300 µg peptides was loaded onto the column in solvent A (5 mM Tris/HCl pH 8.5) and was eluted at a flow rate of 0.25 ml/min using solvent B (5 mM Tris/HCl pH 8.5, 1 M NaCl) in a two-step gradient (0-3 min 0 % B, 3-27 min 27 % B, 27-40 min 100 % B). Fraction collection was initiated after two minutes and continued for 36 minutes. The UV-trace was used to estimate the amount of peptides and neighbouring fractions with low intensity were pooled, resulting in 24 fractions. Fractions were acidified by adding 0.5 % FA and were transferred to pre-equilibrated micro-columns for desalting as previously described (Rappsilber et al., 2007).

#### LC-MS/MS

Peptides were reconstituted in 0.1 % FA and phosphopeptides were reconstituted in 0.1 % FA and 50 mM citric acid, in order to chelate residual Fe^3+^-ions. LC-MS/MS measurements were performed by coupling an Eksigent NanoLC-Ultra 1D+ to a LTQ Orbitrap Velos and a Q-Exactive plus mass spectrometer. Peptides were delivered to a trap column for 10 min at a flow rate of 5 µl/min in loading solvent A (0.1 % FA in water) and were then transferred to the analytical column using a 110 min gradient. Peptides were measured on the LTQ Orbitrap Velos at a flow rate of 300 nl/min and a gradient of 0 - 2 min: 2 - 4 % B; 102 min: 32 % B; 103 min - 106 min: 80 % B, 107-110 min: 2 % B (solvent A: 0.1 % FA, 5 % DMSO; solvent B: 0.1 % FA, 5 % DMSO in ACN). Phosphopeptides were measured on a Q-Exactive plus and a slightly different gradient of 2 min: 2 % B; 98 min: 27 % B; 99 min - 103 min: 80% B, 105-110 min: 2 % B. Both mass spectrometers were operated in data-dependent mode, automatically switching between MS1 and MS2. Full-scan MS1 spectra were acquired at 360 to 1300 m/z. The LTQ Orbitrap Velos was operated at 30,000 resolution, automatic gain control (AGC) target value of 1 x 10^6^ charges and a maximum injection time of 100 ms. Up to 10 precursor ions were allowed for fragmentation. MS2 spectra were acquired at 100 to 2000 m/z at 7,500 resolution, AGC target value of 3 ×10^4^ charges and maximum injection time of 200 ms and ion isolation width was set to 2 Th. The Q-Exactive plus was operated at 70,000 resolution, automatic gain control (AGC) target value of 3 x 10^6^ charges and a maximum injection time of 100 ms. We allowed up to 20 precursor ions to be selected for fragmentation. MS2 spectra were acquired at 200 to 2,000 m/z at 17,500 resolution, AGC target value of 1 ×10^5^ charges and maximum injection time of 50 ms. Precursor ion isolation width was fixed at 1.7 Th and dynamic exclusion was set to 20 s.

#### Data analysis

For data analysis the MaxQuant (Cox and Mann, 2008) (version 1.4.0.5) together with its integrated search engine Andromeda (Cox et al., 2011) was used for the analysis of LC-MS/MS data. Raw files were searched against the UniProtKB database (v.22.07.13, containing 88,381 entries). Carbamidomethylated cysteine was set as a fixed modification. For the phosphoproteome analysis, phosphorylation of serine, threonine and tyrosine, as well as oxidation of methionine and N-terminal protein acetylation were allowed as variable modifications. For raw files of full proteome fractions, only oxidation of methionine and N-terminal protein acetylation were allowed as variable modifications. Enzyme specificity was set to trypsin/P, allowing for cleavage after proline to account for in-source fragmentation. The minimum peptide length was set to seven amino acids and a maximum of two missed-cleavages were allowed. The mass tolerance was set to 4.5 ppm for precursor ions and to 20 ppm for fragment ions. The dataset was adjusted to 1% FDR on the level of proteins and peptide spectrum matches (PSMs).

### Bioinformatics procedures

#### SOM analysis

The self-organizing maps approach is based on a machine-learning algorithm (Wirth et al., 2011; Wirth et al., 2012). It clusters the genes into a grid of gene-groups with similar expression. These gene-groups are called meta-genes and we obtained in an analysis one map per sample/condition. In these maps, the meta-genes are always at the same position, i.e. a meta-gene at coordinates (x,y) for the condition 4-OHTvsEtOH/DMSO 4h contains the same genes as in the 4-OHTvsEtOH/DMSO 72h condition within the same dataset (the meta-genes coordinates are labelled with numbers within the maps). Therefore, the different conditions are directly comparable in terms of gene expression. For further analysis and functional enrichment, clusters of meta-genes (spots) were selected with either high or low expression compared to the overall expression range of the condition. The high expression spots must have an expression range between the maximal expression and 65% of the maximal expression according to (Wirth et al., 2011; Wirth et al., 2012). Same for the low expression spots with the minimal expression. Due to the varying numbers of genes in the three different datasets, ranging from 26,000 at mRNA level to 500 at the phosphoprotein level, we decided to use a grid of 10x10 meta-genes, however, all measured genes were used for the analyses. We decided to use log2FCs between 4-OHT and EtOH/DMSO to perform the SOM analyses for all the datasets.

#### MicroRNA target prediction

To discover potential miRNA targets, we used the miRTarBase v6.1 (Chou et al., 2018) as target database and a modified version of the miRlastic R package (Sass et al., 2015). The miRlastic approach uses a linear regression model with elastic net regularization to predict the miRNA targets based on the miRNA and mRNA expression. The elastic net regression is penalized to fulfil the known anti-correlation between miRNA and mRNA (Guo et al., 2010). Differentially expressed miRNAs as well as the differentially expressed mRNAs for each time point were used. For the miRNA target prediction, the log2FC between the 4-OHT (mutant HRAS induced) and the EtOH/DMSO control condition for every time point are used. All miRNAs with an adjusted p-value below 0.1 as well as all mRNAs with an adjusted p-value of 0.1 were considered.

#### Transcription factor identification

To identify differentially expressed transcription factors among the proteome, phosphoproteome and transcriptome datasets, we searched for the corresponding annotations in the Animal TFDB 2.0 (Zhang et al., 2015). We prepared a unified list based on proteome and phosphoproteome data. We annotated the transcription factors separately in transcriptome datasets obtained at the four time points following induction of HRAS^G12V^. To identify the common transcription factors for each time point, the overlap of the proteomic transcription factors and the transcriptomic transcription factors was determined. The resulting four lists of common transcription factors were subsequently integrated to build a time-independent list. With this procedure, we identified 27 common transcription factors deregulated at the different molecular levels.

